# Zika virus, a new threat for Europe?

**DOI:** 10.1101/048454

**Authors:** Henri Jupille, Goncalo Seixas, Laurence Mousson, Carla A. Sousa, Anna-Bella Failloux

## Abstract

**Background:** Since its emergence in 2007 in Micronesia and Polynesia, the arthropod-borne flavivirus Zika virus (ZIKV) has spread in the Americas and the Caribbean, following first detection in Brazil in May 2015. The risk of ZIKV emergence in Europe increases as imported cases are repeatedly reported. Together with chikungunya virus (CHIKV) and dengue virus (DENV), ZIKV is transmitted by *Aedes* mosquitoes. Any countries where these mosquitoes are present could be potential sites for future ZIKV outbreak.

**Methodology/Principal Findings:** Mosquito females were challenged with an Asian genotype of ZIKV. Fully engorged mosquitoes were then maintained in insectary conditions (28°±1°C, 16h:8h light:dark cycle and 80% humidity). 16-24 mosquitoes from each population were examined at 3, 6, 9 and 14 days postinfection to estimate the infection, disseminated infection and transmission rates. Based on these experimental infections, we demonstrated that *Ae. albopictus* from France were not very susceptible to ZIKV.

**Conclusions/Significance:** In combination with the restricted distribution and lower population densities of European *Ae. albopictus*, our results corroborate the low risk for ZIKV to expand into most parts of Europe with the possible exception of the warmest regions bordering the Mediterranean coastline.

**Author summary:** In May 2015, local transmission of Zika virus (ZIKV) was reported in Brazil and since then, more than 1.5 million human cases have been reported in Latin America and the Caribbean. This arbovirus, primarily found in Africa and Asia, is mainly transmitted by *Aedes* mosquitoes, *Aedes aegypti* and *Aedes albopictus*. Viremic travelers returning from America to European countries where *Ae. albopictus* is established can become the source for local transmission of ZIKV. In order to estimate the risk of seeding ZIKV into local mosquito populations, the ability of European *Ae. aegypti* and *Ae. albopictus* to transmit ZIKV was measured using experimental infections. We demonstrated that *Ae. albopictus* and *Ae. aegypti* from Europe were not very susceptible to ZIKV. The threat for a Zika outbreak in Europe should be limited.

## Introduction

Zika virus (ZIKV) (genus *Flavivirus*, family *Flaviviridae)* is an emerging arthropod-borne virus transmitted to humans by *Aedes* mosquitoes. ZIKV infection in humans was first observed in Africa in 1952 [1], and can cause a broad range of clinical symptoms presenting as a “dengue-like” syndrome: headache, rash, fever, and arthralgia. In 2007, an outbreak of ZIKV on Yap Island resulted in 73% of the total population becoming infected [2]. Following this, ZIKV continued to spread rapidly with outbreaks in French Polynesia in October 2013 [3], New Caledonia in 2015 [4], and subsequently, Brazil in May 2015 [5, 6]. During this expansion period, the primary transmission vector is considered to have been *Aedes aegypti*, although *Aedes albopictus* could potentially serve as a secondary transmission vector [7]. As Musso et al. [8] observed, the pattern of ZIKV emergence from Africa, throughout Asia, to its subsequent arrival in South America and the Caribbean closely resembles the emergence of Chikungunya virus (CHIKV). In Europe, returning ZIKV-viremic travelers may become a source of local transmission in the presence of *Aedes* mosquitoes, *Ae. albopictus* in Continental Europe and *Ae. aegypti* in the Portuguese island of Madeira. *Ae. albopictus* originated from Asia and was recorded for the first time in Europe in Albania in 1979 [9], then in Italy in 1990 [10]. It is now present in all European countries around the Mediterranean Sea [11]. This mosquito was implicated as a vector of CHIKV and DENV in Europe [12]. On the other hand, *Ae. aegypti* disappeared after the 1950s with the improvement of hygiene and anti-malaria vector control. This mosquito reinvaded European territory, Madeira island, in 2005 [13], and around the Black Sea in southern Russia, Abkhazia, and Georgia in 2004 [11]. The species was responsible for outbreaks of yellow fever in Italy in 1804 [14] and dengue in Greece in 1927–1928 [15]. To assess the possible risk of ZIKV transmission in Europe, we compared the relative vector competence of European *Ae. aegypti* and *Ae. albopictus* populations to the Asian genotype of ZIKV.

## Materials and Methods

### Ethics Statement

The Institut Pasteur animal facility has received accreditation from the French Ministry of Agriculture to perform experiments on live animals in compliance with the French and European regulations on care and protection of laboratory animals. This study was approved by the Institutional Animal Care and Use Committee (IACUC) at the Institut Pasteur. No specific permits were required for the described field studies in locations that are not protected in any way and did not involve endangered or protected species.

### Mosquitoes

Four populations of mosquitoes (two populations of *Ae. aegypti:* Funchal and Paul do Mar, collected on island of Madeira and two populations of *Ae. albopictus:* Nice and Bar-sur-Loup in France) were collected using ovitraps. Eggs were immersed in dechlorinated tap water for hatching. Larvae were distributed in pans of 150-200 individuals and supplied with 1 yeast tablet dissolved in 1L of water every 48 hours. All immature stages were maintained at 28°C ± 1°C.

After emergence, adults were given free access to a 10% sucrose solution and maintained at 28°C ± 1°C with 70% relative humidity and a 16:8 light/dark cycle. The F1 generation of *Ae. aegypti* from Madeira and F7-8 generation of *Ae. albopictus* from France were used for experimental infections.

### Viral strain

The ZIKV strain (NC-2014-5132) originally isolated from a patient in April 2014 in New Caledonia was used to infect mosquitoes. The viral stock used was subcultured five times on Vero cells prior to the infectious blood-meal. The NC-2014-5132 strain is phylogenetically closely related to the ZIKV strains circulating in the South Pacific region, Brazil [5] and French Guiana [16].

### Oral Infection of Mosquitoes

Infectious blood-meals were provided using a titer of 10^7^ TCID_50/_mL. Seven-day old mosquitoes were fed on blood-meals containing two parts washed rabbit erythrocytes to one part viral suspension supplemented with ATP at a final concentration of 5 mM. Engorged females were transferred to cardboard containers with free access to 10% sucrose solution and maintained at 28°C and 70% relative humidity with a 16:8 light/dark cycle. 16-24 female mosquitoes from each population were analyzed at 3, 6, 9, and 14 days post-infection (dpi) to estimate the infection, disseminated infection and transmission rates. Briefly, legs and wings were removed from each mosquito followed by insertion of the proboscis into a 20 μL tip containing 5 μL FBS for 20 minutes. The saliva-containing FBS was expelled into 45 μL serum free L-15 media (Gibco), and stored at ‐80°C. Following salivation, mosquitoes were decapitated and head and body (thorax and abdomen) were homogenized separately in 300 L-15 media supplemented with 3% FBS using a Precellys homogenizer (Bertin Technologies) then stored at ‐80°C. Infection rate was measured as the percentage of mosquitoes with infected bodies among the total number of analyzed mosquitoes. Disseminated infection rate was estimated as the percentage of mosquitoes with infected heads (i.e., the virus had successfully crossed the midgut barrier to reach the mosquito hemocoel) among the total number of mosquitoes with infected bodies. Transmission rate was calculated as the overall proportion of females with infectious saliva among those with disseminated infection. Samples were titrated by plaque assay in Vero cells.

### Virus Quantification

For head/body homogenates and saliva samples, Vero E6 cell monolayers were inoculated with serial 10-fold dilutions of virus-containing samples and incubated for 1 hour at 37°C followed by an overlay consisting of DMEM 2X, 2% FBS, antibiotics and 1% agarose. At 7 dpi, overlay was removed and cells were fixed with crystal violet (0.2% Crystal Violet, 10% Formaldehyde, 20% ethanol) and positive/negative screening was performed for cytopathic effect (body homogenates) or plaques were enumerated (head homogenates and saliva samples). Vero E6 cells (ATCC CRL-1586) were maintained in DMEM (Gibco) supplemented with 10% fetal bovine serum (Eurobio), Penicillin and Streptomycin, and 0.29 mg/mL l-glutamine.

### Statistical analysis

All statistical tests were conducted using the STATA software (StataCorp LP, Texas, USA) using Fisher’s exact test and P-values>0.05 were considered non-significant.

## Results

### Aedes aegypti from Madeira transmit ZIKV efficiently

To test whether *Ae. aegypti* from a European territory were able to transmit ZIKV, we analyzed the vector competence of two *Ae. aegypti* populations collected on the island of Madeira based on three parameters: viral infection of the mosquito midgut, viral dissemination to secondary organs, and transmission potential, analyzed at 3, 6, 9, and 14 dpi. The two populations presented similar infection and disseminated infection (Figure) (P > 0.05) with the highest rates measured at 9 dpi and 9-14 dpi, respectively. Only mosquitoes presenting an infection (i.e. infected midgut) were analyzed for viral dissemination and only mosquitoes with a disseminated infection were assessed for transmission success. Thus, only *Ae. aegypti* Funchal were able to transmit ZIKV at 9 and 14 dpi (Figure).

**Figure.**
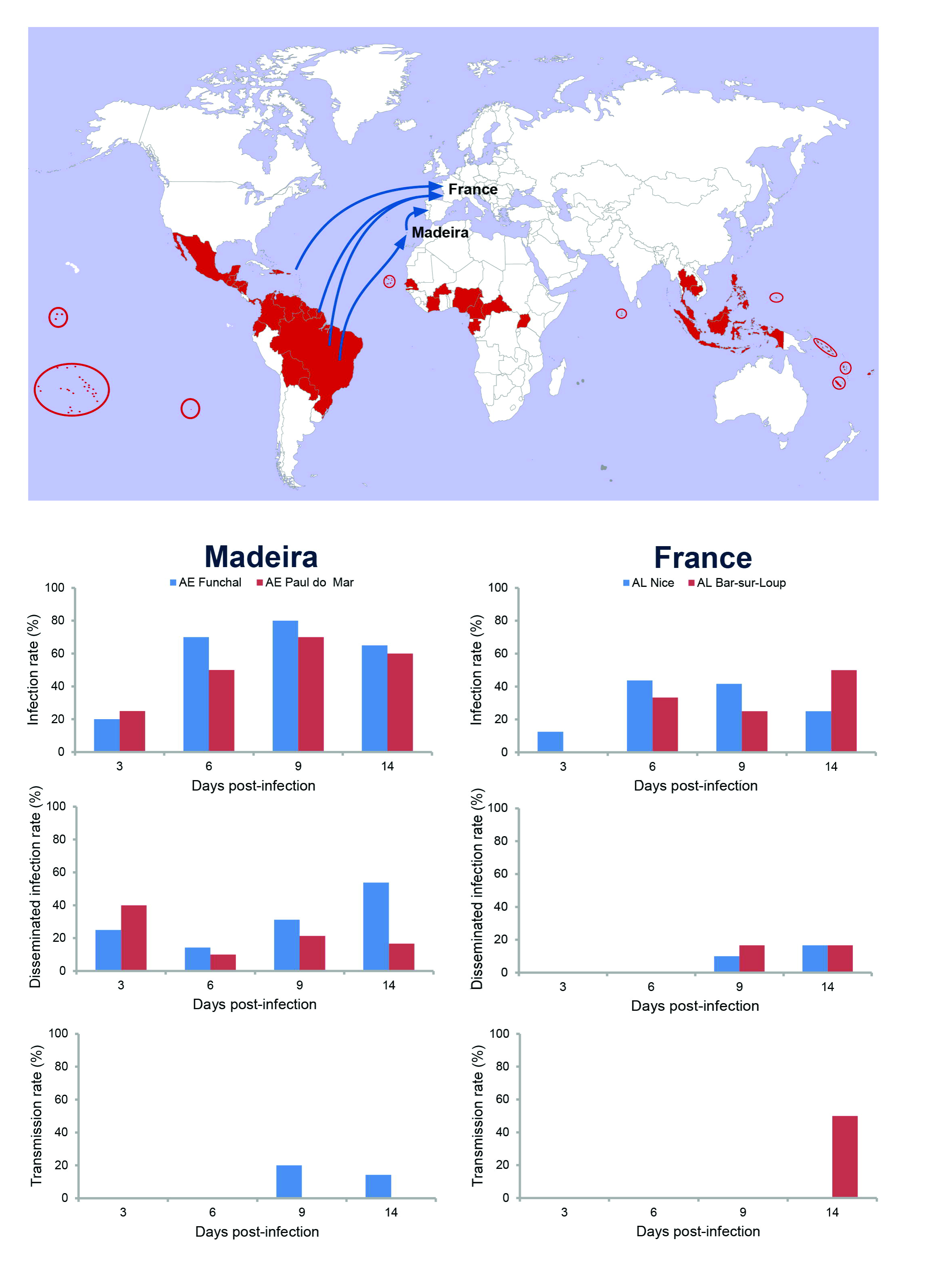
*Ae. aegypti* from Madeira Island and *Ae. albopictus* from France were assessed for viral infection, dissemination, and transmission at days 3, 6, 9, 14 after infection with ZIKV provided at a titer of 10^7^ TCID_50_/mL. 16-24 mosquitoes were sampled each day. Infection rates were measured as the percentage of mosquitoes with infected bodies among the total number of analyzed mosquitoes. Disseminated infection rates were estimated as the percentage of mosquitoes with infected heads (i.e., the virus has successfully crossed the midgut barrier to reach the hemocoel) among the total number of mosquitoes with infected bodies. The transmission rate was calculated as the overall proportion of females with infectious saliva among those with disseminated ZIKV infection. AE = *Ae. aegypti*; AL = *Ae. albopictus*. In red, countries where ZIKV has been isolated.

### French Ae. *albopictus* showed significantly reduced competence to transmit ZIKV

To determine if *Ae. albopictus* present in continental Europe were able to sustain local transmission of ZIKV as previously observed with CHIKV and DENV, we evaluated the vector competence of two *Ae. albopictus* populations collected in Nice and Bar-sur-Loup in the South of France. When compared with *Ae. aegypti*, the two *Ae. albopictus* populations showed equivalent but reduced infection and disseminated infection (Figure) (P > 0.05) with highest rates observed at 6 dpi and 14 dpi, respectively. Only one individual among two *Ae. albopictus* Barsur-Loup having ensured viral dissemination was able to transmit ZIKV at 14 dpi (Figure).

In summary, virus titers measured in *Ae. albopictus* were much lower than those detected in *Ae. aegypti*. Virus dissemination through *Ae. aegypti* was noticeably superior and consequently, viral loads in saliva were higher for *Ae. aegypti* (1500 pfu compared with 2 pfu at 9 dpi; data not shown). Moreover, transmission of ZIKV occurred earlier and in a much higher proportion of *Ae. aegypti* when compared with *Ae. albopictus*.

## Discussion

ZIKV could be transmitted, spread and maintained in Europe either via (i) Madeira where the main vector *Ae. aegypti* has been established since 2005 or (ii) Continental Europe where *Ae. albopictus* is known to have been present since 1979 [11]. We demonstrated that ZIKV was amplified and transmitted efficiently by European *Ae. aegypti* from Madeira. This contrasts with the much lower vector competence for ZIKV amplification and transmission of French *Ae. albopictus*. Taking these observations and the overall average lower temperatures of most regions of Europe into account, the risk of major outbreaks of Zika fever in most areas of Europe, at least for the immediate future, appears to be relatively low.

Our results highlight the potential risk for ZIKV transmission on Madeira where two main factors are present: the presence of the main vector, *Ae. aegypti* introduced in 2005 [17] and imported cases from Brazil with which Madeira, an autonomous region of Portugal, maintains active exchanges of goods and people sharing the same language. Thus Madeira Island could be considered as a stepping stone for an introduction of ZIKV into Europe.

Autochthonous cases of CHIKV and DENV have been reported in Europe since 2007: CHIKV in Italy in 2007, South France in 2010, 2014, and DENV in South France in 2010, 2013, 2015, and Croatia in 2010 [18]. The invasive species *Ae. albopictus* first detected in Europe in 1979 [9] has played a central role in this transmission [18]. Thus, there might be a risk of a similar establishment of ZIKV in Europe upon the return of viremic travelers [19, 20]. We showed that *Ae. albopictus* from South France were less competent for ZIKV infection requiring 14 days to be excreted in the mosquito saliva after infection. Therefore, we can suggest that the Asian tiger mosquito from Southern France and more widely, Europe, are less suitable to sustain local transmission of ZIKV compared to CHIKV and perhaps, DENV.

Considering the extensive airline travel between Latin America and Europe, the risk for local transmission of ZIKV in the European area where the mosquito *Ae. albopictus* is widely distributed, is assumed to be minimal based on our studies of vector competence. Nevertheless, reinforcement of surveillance and control of mosquitoes should remain a strong priority in Europe since *Aedes* mosquitoes also transmit DENV and CHIKV and virus adaptation to new vectors cannot be excluded, as previously observed with CHIKV in La Reunion [21, 22].

## Acknowledgments

The authors thank Myrielle Dupont-Rouzeyrol for providing the ZIKV strain. We are grateful to Marie Vazeille for technical assistance in performing mosquito titrations. We warmly thank Xavier de Lamballerie, Marc Lecuit, Anavaj Sakuntabhai, Richard Paul and Alain Kohl for their support. We also thank Pascal Delauney, Christophe Lagneau and Gregory Lambert for mosquito collections in South of France. We deeply thank Ana Clara Silva for the work developed in Madeira.

## References

1. Dick GW, Kitchen SF, Haddow AJ. Zika virus. I. Isolations and serological specificity. Trans R Soc Trop Med Hyg 1952; 46:509–20.

2. Duffy MR, Chen TH, Hancock WT, et al. Zika virus outbreak on Yap Island, Federated States of Micronesia. N Engl J Med 2009; 360:2536–43.

3. Cao-Lormeau VM, Roche C, Teissier A, et al. Zika virus, French Polynesia, South pacific, 2013. Emerg Infect Dis 2014; 20:1085–6.

4. Dupont-Rouzeyrol M, O'Connor O, Calvez E, et al. Co-infection with Zika and dengue viruses in 2 patients, New Caledonia, 2014. Emerg Infect Dis 2015; 21:381–2.

5. Zanluca C, de Melo VC, Mosimann AL, Dos Santos GI, Dos Santos CN, Luz K. First report of autochthonous transmission of Zika virus in Brazil. Mem Inst Oswaldo Cruz 2015; 110:569–72.

6. Campos GS, Bandeira AC, Sardi SI. Zika Virus Outbreak, Bahia, Brazil. Emerg Infect Dis 2015; 21:1885–6.

7. Wong PS, Li MZ, Chong CS, Ng LC, Tan CH. Aedes (Stegomyia) albopictus (Skuse): a potential vector of Zika virus in Singapore. PLoS Negl Trop Dis 2013; 7:e2348.

8. Musso D, Cao-Lormeau VM, Gubler DJ. Zika virus: following the path of dengue and Chikungunya? Lancet 2015; 386:243–4.

9. Adhami J, Reiter P. Introduction and establishment of Aedes (Stegomyia) albopictus skuse (Diptera: Culicidae) in Albania. J Am Mosq Control Assoc 1998; 14:340–3.

10. Sabatini A, Raineri V, Trovato G, Coluzzi M. [Aedes albopictus in Italy and possible diffusion of the species into the Mediterranean area]. Parassitologia 1990; 32:301–4.

11. Medlock JM, Hansford KM, Schaffner F, et al. A review of the invasive mosquitoes in Europe: ecology, public health risks, and control options. Vector Borne Zoonotic Dis 2012; 12:435–47.

12. Vega-Rua A, Zouache K, Caro V, et al. High efficiency of temperate Aedes albopictus to transmit chikungunya and dengue viruses in the Southeast of France. PLoS One 2013; 8:e59716.

13. Nene V, Wortman JR, Lawson D, et al. Genome sequence of Aedes aegypti, a major arbovirus vector. Science 2007; 316:1718–23.

14. Levre E. The yellow fever outbreak of 1804 in Leghorn. Ann Ig 2002; 14:153–7.

15. Rosen L. Dengue in Greece in 1927 and 1928 and the pathogenesis of dengue hemorrhagic fever: new data and a different conclusion. Am J Trop Med Hyg 1986; 35:642–53.

16. Antoine Enfissi JC, Jimmy Roosblad, Mirdad Kazanji, Dominique Rousset Zika virus genome from the Americas. The Lancet 2015; 7 janvier 2016.

17. Margarita Y SGA, Lencastre I, Silva AC, Novo T, Sousa C, et al. First record of Aedes (Stegomia) aegypti (Linnaeus, 1762) (Diptera, Culicidae) in Madeira Island - Portugal. Acta Parasitológica Portuguesa 2006; 13:59–61.

18. Rezza G. Dengue and chikungunya: long-distance spread and outbreaks in naive areas. Pathog Glob Health 2014; 108:349–55.

19. Tappe D, Rissland J, Gabriel M, et al. First case of laboratory-confirmed Zika virus infection imported into Europe, November 2013. Euro Surveill 2014; 19.

20. Zammarchi L, Tappe D, Fortuna C, et al. Zika virus infection in a traveller returning to Europe from Brazil, March 2015. Euro Surveill 2015; 20.

21. Tsetsarkin KA, Vanlandingham DL, McGee CE, Higgs S. A single mutation in chikungunya virus affects vector specificity and epidemic potential. PLoS Pathog 2007; 3:e201.

22. Vazeille M, Moutailler S, Coudrier D, et al. Two Chikungunya isolates from the outbreak of La Reunion (Indian Ocean) exhibit different patterns of infection in the mosquito, Aedes albopictus. PLoS One 2007; 2:e1168.

